# Culturomics from field-grown crop plants using dilution-to-extinction, two-step library preparation and amplicon sequencing

**DOI:** 10.1101/2025.03.04.641206

**Authors:** Eglantina Lopez-Echartea, Nicholas Dusek, Mallory Misialek, Mohammad Al Mahmud Un Nabi, Riley Williamson, Komal Marathe, Barney A. Geddes

**Author notes:** Corresponding author and email address: Corresponding author: Barney Geddes. Repositories: Sequencing data was deposited in the NCBI SRA under the BioProject PRJNA1220177. R scripts used for data analysis are available at https://github.com/NDSU-Geddes-Lab/cultured-microbe-identification. Protocols for automated library preparation have been published on protocols.io and are available at DOI: dx.doi.org/10.17504/protocols.io.eq2ly662egx9/v1.

## Abstract

Culturomics approaches have advanced microbial research by enabling the high-throughput isolation and characterization of a broader range of bacterial taxa, including some previously considered unculturable. Here, we present the testing and optimization of a protocol for isolating and identifying hundreds of cultivable microbes from field-grown plants. This protocol was tested and optimized using the root microbiomes of field-grown corn and pea plants under varying environmental conditions in North Dakota, USA. By employing dilution-to-extinction culturing and a two-step barcoding PCR strategy targeting the V4 region of the 16S rRNA gene, we identified over 200 unique bacterial isolates. The optimized bioinformatic pipeline, built around the DADA2 package, ensured accurate amplicon sequence variant (ASV) detection and taxonomy assignment. The resulting bacterial isolates span diverse phylogenetic groups, including plant-associated taxa known for promoting plant growth and mitigating stress. Our findings highlight the value of culturomics in generating microbial collections for synthetic community design and advancing plant-microbe interaction research. The protocol’s scalability, cost-effectiveness, and robust performance demonstrate its potential for widespread application in agricultural microbiome studies.

**Impact statement:** High-throughput isolation and characterization of cultivable microbes from plant microbiomes is crucial for advancing microbiome research. However, efficiently recovering a diverse range of bacterial taxa remains a challenge due to high costs and labor-intensive protocols. Our optimized culturomics protocol integrates dilution-to-extinction culturing with a two-step barcoding PCR strategy, enhancing recovery rates and reducing costs while maintaining high accuracy. By employing next-generation sequencing (NGS) and a streamlined bioinformatic pipeline built around the robust DADA2 workflow for amplicon sequencing, the method enables the scalable recovery of hundreds of unique bacterial isolates. This approach makes significant advancements over traditional culturing methods and other high-throughput cultivation protocols, providing an efficient, cost-effective platform for generating microbial collections essential for synthetic communities, comparative genomics, and agricultural microbiome applications.

**Data summary:** All protocols, sequence data, and analysis codes have been made publicly available in open-access repositories. The authors confirm all supporting data, code, and protocols have been provided within the article or through supplementary data files.

## 5. Introduction

The first isolates of pure bacterial cultures were introduced in the late 19th century by traditional solid culture media^1^. Although this technique has been one of the pillars of microbiology as research field, advances in culturing microbes have not kept pace in recent years with advances in our ability to characterize microbial communities by next-generation sequencing^2,3^. However, in this multi-omics era where thousands of terabytes of sequence data are generated year after year, cultivation is still irreplaceable.

Microbial cultures are necessary for the evaluation of metagenomics data, the assessment and characterization of genes and genomes, the isolation of bacterial and archaeal viruses and testing hypotheses^4^. Moreover, the characterization and isolation of new bacterial isolates is of high relevance due to their multiple applications in biotechnology, bioremediation, bioactive compounds and bioprospection^5–9^. The field of agriculture will also benefit from the high-throughput cultivation and fast identification of microbes as the plant microbiome could provide protection against pathogens, fix nitrogen, promote plant growth and decrease stress from abiotic factors such as drought, salinity and contaminants^10–14^. Combining cultured microbes into synthetic bacterial communities (SynComs) has also gained significant traction in recent years to facilitate a reductionist, “bottom-up” approach to study of microbial ecology and complement the wealth of “top-down” amplicon sequencing studies^15–17^.

Microbes represent the largest set of biomass on Earth and are represented in all domains of life, yet 90% of their diversity is still unexplored^18^, in part because most of the prokaryotic taxa have not been yet cultured ^4^. The reason for this uncultivability relies on several factors such as, the absence of factors produced by other microbes^19,20^, interspecies interactions such as symbiotic growth^21,22^, and variations in the growth rate^23^ and dormancy^24^ of individual microbes. Several groups have attempted to increase culturability through incorporating radically altered approaches and technologies such as microfluidic droplets^25^, microcapsules^26^ and miniature diffusion chambers that are placed in the environment^27^. Still, perhaps the most significant advances in microbial culturing have resulted from more incremental improvements on traditional culturing methods by optimizing culture conditions. For example, the improvement in media, nutrient addition, and approaches like ‘dilution-to-extinction’ have made great advances in fields like marine microbiology where microbes are slow growers with low nutrient requirements^3,28,29^. Pioneered in the human gut microbiome, cutting-edge approaches dubbed “culturomics” involve combining optimized growth conditions with high-throughput workflows enabled by robotics, automation and next-generation sequencing approaches for identification of target microbes of interest^30–33^.

Culturomics approaches have been applied to the plant microbiome and shown great success where the proportion of cultivatable taxa has been found to be relatively high^34,35^. In this study, we further advance culturomics efforts in the plant microbiome, by 1) testing the dilution-to-extinction approach for cultivation of microbes from field-grown corn and pea plants in North Dakota, USA, and 2) developing a two-step PCR and bioinformatic strategy for identification of cultured microbes. We combined these tests into an updated protocol for culturomics from the plant microbiome.

## 6. Theory and implementation

### Methods

#### Sampling and sample preparation

Pea plants were collected in July 2022 from a Fusarium/Aphanomyces root rot trial in Williston, North Dakota and a salinity trial in Carrington, North Dakota. Corn plants were collected in July 2023 from a field trial in Gardner, North Dakota that included variable nitrogen fertilizer application rates (0 or 200 lbs/ac) at planting. Samples were collected from field locations, leaving the rhizosphere intact and transported on ice to the laboratory. Samples were processed in the laboratory by removing loosely associated microbes with a series of washes in phosphate-buffered saline, followed by cutting and grinding of plant tissues into a slurry, as previously described^34^. An aliquot of the slurry was separated for microbial community composition analysis and the remaining was used for culturing.

#### Bacterial culturing

A slurry from each plant root sample was used for culturing by dilution-to-extinction in 10% tryptic soy broth (TSB) as previously described^34^, and included a 3-fold serial dilution from an initial 2000x dilution of the plant slurry to 486,000x. A total of 20 plates for each dilution were inoculated for each plant sample. Culture plates were incubated for 12 days at room temperature. Individual 96-well plates from dilution-to-extinction culturing were then selected based on the proportion of growth in wells (18-55%). An aliquot from each well was transferred to a new 96-well plate for cultured bacteria identification, and the remaining contents were combined with glycerol (40% final volume) and stored at -80°C.

For purification of isolates, streak-plating from single colonies was used on TSB agar plates. Following three rounds of purification, isolates were resuspended in TSB and combined with glycerol (40% final volume) for long-term storage.

#### PCR amplification for bacterial identification

Genomic DNA was extracted from bacterial culture aliquots using an alkaline lysis method. Amplification of the bacterial 16S rRNA gene was performed using an adaptation to the standard Illumina two-step PCR approach^36^. Specifically, we incorporated 4-6 nucleotide barcodes into twelve unique forward and eight unique reverse primers targeting the V4 region of the 16S rRNA gene (V4_515F and V4_R) in between the V4 binding site and adaptor tail binding sites for secondary amplification (**Supplementary Table 2**). These primers were used with a unique forward primer for each column of the 96-well plate and each reverse primer for each row such that every well would receive a unique primer combination. The primary PCR amplification used KAPA HotStart polymerase (Kapa Biosystems, USA), with cycling conditions of an initial denaturation at 95°C for 3 minutes, followed by 26 cycles of 95°C for 30 seconds, 55°C for 30 seconds, and 72°C for 30 seconds, with a final extension at 72°C for 5 minutes. PCR products from each of the 96 wells were then pooled together and purified using Mag-Bind beads (Omega Bio-tek, USA). A second round of PCR was performed on each pooled plate sample from PCR1, incorporating Nextera primers with sequencing adapters^34^, under similar conditions but reduced to 9 cycles. The PCR products were again purified using Mag-Bind beads (Omega Bio-tek, USA), followed by quantification using a Quant-iT PicoGreen dsDNA Assay Kit (Thermo Fisher Scientific, USA). Libraries were pooled in equimolar amounts and sequenced on an Illumina MiSeq platform on the Core Lab of the Department of Microbiological Sciences of the North Dakota State University following standard protocols.

PCR for root slurry microbiome profiling was performed using the same reagents and cycling conditions as described for bacterial isolate amplification. The only difference was that the PCR1 primers (V4_515F and V4_R, **Supplementary Table 2**) did not include the 4-6 nucleotide barcodes used in the high-throughput cultivation workflow.

PCR amplification and Sanger sequencing were used to validate the bacterial 16S rRNA gene sequences from purified isolates. The full-length 16S rRNA gene was amplified using 27F and 1492R primers, following previous PCR1 conditions but with an extended annealing time of 1 minute. PCR products were treated with ExoSAP-IT (Thermo Fisher Scientific, USA) and sequenced by Molecular Cloning Laboratories, USA using Sanger sequencing.

#### Bioinformatic analysis

FASTQ files from Illumina sequencing from the cultivation plates were demultiplexed using a custom script (microbeID.R from NDSU-Geddes-Lab/cultured-microbe-identification). Briefly, the script trims, merges and denoises reads using the DADA2 package^37^, demultiplexes sequences from each plate and assigns them to wells based on the primer barcode combination and reported read number, purity and taxonomy of amplicon sequence variants in each well. The FASTQ files from Illumina sequencing from the root slurry were processed in the same way as the in Benz et al., 2024 ^38^.

For the construction of the phylogenetic treess V4 16S rRNA gene DNA sequences in FASTA format from the root slurry and the cultivation plates were aligned using multiple sequence alignment in R with the DECIPHER^39^ and phangorn^40^ packages. Briefly, the aligned sequences were used to compute a distance matrix using maximum likelihood, and an initial Neighbor-Joining tree was constructed. The tree was further refined using a maximum likelihood approach by fitting a general time reversible model. The optimized phylogenetic tree was saved as a newick file and further processed in iTOL^41^ to visualize and label the trees effectively.

### Results and Discussion

#### Microbe isolation from field-grown crop plants by dilution-to-extinction

We set out to test high throughput culturomics in our experimental systems (field-grown crop plants with biotic and abiotic stress), starting with a previously published protocol that optimized the approach for root microbiome culturing from the plants^34^. Dilution-to-extinction relies on achieving a concentration where individual wells predominantly contain single cells, minimizing the competition between microbes and allowing for growth of slower-growing or nutrient-limited species. This is done by selecting specific dilutions with a proportional growth in wells that indicates most wells have been derived from a single viable bacterial cell according to a Poisson distribution model^34^. We tested a range of dilutions, from 2,000x to 486,000x, across different plant root samples (pea and corn) and environmental conditions (disease, salinity, fertilized, non-fertilized) collected from the field. Successful dilutions were identified based on growth rates in 96-well plates, with ideal dilutions defined as those yielding ∼35% growth. At these levels, there is a high likelihood that each well contains either no cells or only a single viable cell.

For pea plants, we selected one dilution that closely matched these criteria (33% growth in the 54,000x dilution for peas growing in the disease inoculated fields and 38% growth in the 18,000x dilution for peas growing in the salinity induced fields). For corn plants, we selected the following dilutions: 18,000x dilution (55% growth) for corn plants growing in 200 lbs/acre N fertilizer field and 6,000x and 18,000x dilutions (30% and 21% growth, respectively) for corn plants growing in the non-fertilized fields (**Table 1**). Overall, the dilutions within the range of the series recommended by Zhang et al. (up to 54,000x) were successful for our conditions, though given that in one case^34^, only the maximum dilution was successful. Including further dilutions to 486,000X may be important for some plants with dense microbiome colonization such as those grown in rich field soil.

**Table 1.** Proportion of well growth from dilution-to-extinction.

| dilution | Pea (disease) | Pea (salinity) | Corn (0) | Corn(200) |
| --- | --- | --- | --- | --- |
| 2000x | TM | TM | TM | TM |
| 6,000x | TM | TM | 30% | TM |
| 18,000x | TM | 38% | 21% | 55% |
| 54,000x | 33% | TF | TF | TF |
| 162,000x | TF | TF | TF | TF |
| 486,000x | TF | TF | TF | TF |

TM= Too many >60% growth
TF= Too few <20% growth

#### Two-step library preparation for identification of cultivated bacteria by amplicon sequencing

We employed barcoded PCR to facilitate the characterization our cultured microbe samples. However, we chose to develop a new barcoding approach relative to the published protocol by Zhang et al. due to a high cost of long primers required for the two-sided barcode approach utilized^34^. We developed an alternate strategy for barcode PCR that utilized two-step library preparation which has been reported has having improved accuracy relative to one-step methods^36^. In two-step PCR we were able to incorporate barcodes into the primary PCR primers for 16S rRNA gene amplification while still keeping the overall length relatively short and thus cost-effective (less than 60 bp). During this step, 96 wells from each plate get barcoded by the combination of twelve different forward and eight different reverse primers (Fig. 2) described in **Supplementary Table 2**. We chose to utilize primers that targeted the V4 region of 16S rRNA gene for consistency with our routine microbiome workflows. The secondary PCR utilized the same primer sets as used for this method in routine microbiome sequencing^34^, and the Illumina Nextera barcodes in secondary PCR primers barcoded each plate after combining primary barcoded wells from the first PCR into a single sample.

#### Bioinformatic pipeline to analyze amplicon sequences from cultivated bacteria

To analyze multiplexed microbiome sequences from 96-well culture plates, we developed a pipeline in R using the DADA2 package^37^, along with some custom functions for primer demultiplexing and sequence purity calculations. The pipeline is designed to be run as a command line script, and performs four basic steps: 1) quality filtering, trimming, and merging of raw forward and reverse sequence reads; 2) assignment of sequences to plate wells based on forward and reverse primer sequences; 3) counting and purity calculation of unique sequences with respect to their plate wells; and 4) taxonomy assignment of unique ASVs. The output of the pipeline is a CSV file containing the ASV, the taxonomy, and the top *n* “hits” (i.e., plate wells, ranked by count) for each unique sequence. For each of the top *n* hits, the plate ID, well ID, sequence count, and sequence purity are reported (**Table 2**).

**Table 2.**
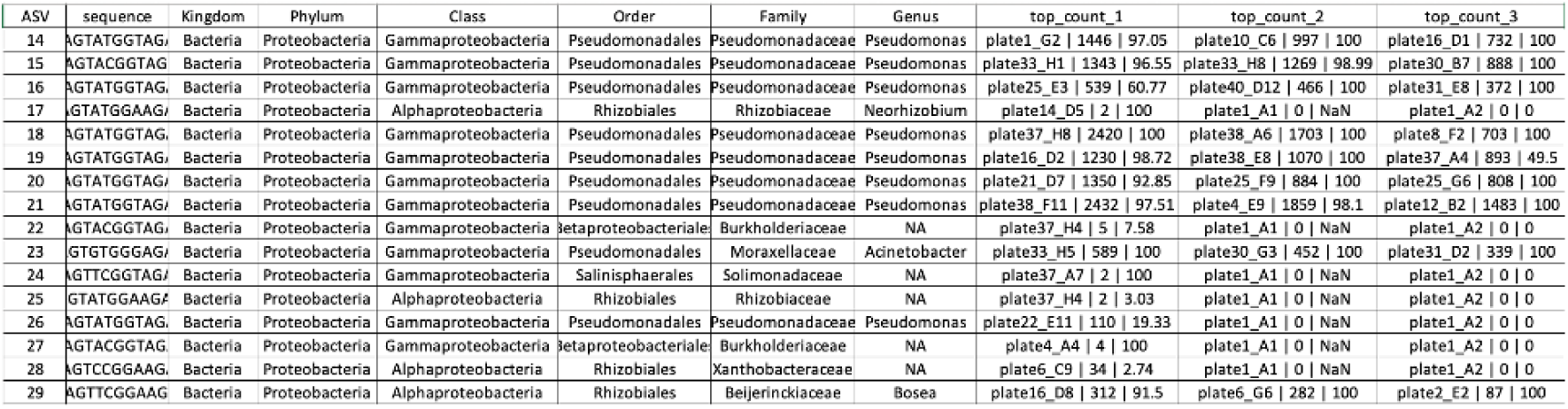
Example output of cultivated bacteria analysis pipeline.

#### Comparisons of cultivated bacteria identity to the root slurry metagenome composition

To investigate culture biases in our approaches, we generated phylogenetic trees from the ASVs identified in the high-troughput cultivation (HTPC) pipeline (∼250 each for pea and corn) and the top 250 ASVs identified by amplicon sequencing of the slurry microbiome used for culturing. Overall, we observed widespread representation of groups of microbes that are associated with plant microbiomes including members of classes Bacteroidia, Bacilli, Actinobacteria, and Alpha-, Beta- and Gamma-proteobacteria. A more even distribution of HTPC and slurry ASVs are seen in the corn microbiome (**Figure 1**), whereas the pea microbiome showed more evidence of bias. Less dominant classes with several representatives in the slurry but missing putatively culturable isolates included Saccharimonadia, Thermoleophilia and Acidmicrobiia. At the order level, most that had several representatives in the slurry included putatively culturable representatives in the pipeline. Orders with several representatives but without putatively culturable isolates included Micromonosporales, Pseudocardiales and Steroidobacterales.

**Figure 1:**
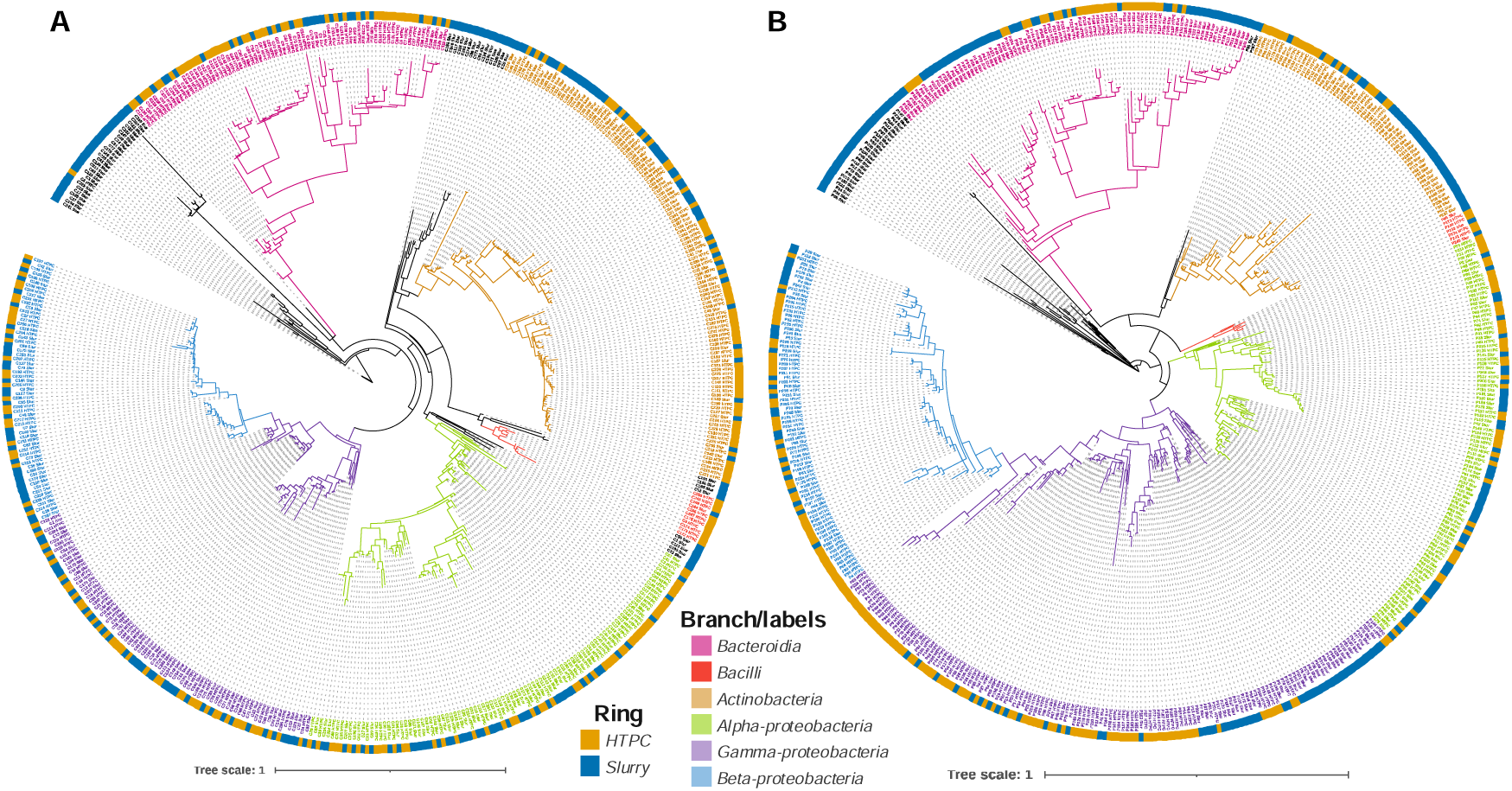
Maximum likelihood phylogenetic trees comparing the position of putatively culturable ASVs from the HTPC pipeline to the top 250 ASVs by abundance from the pea **(A)** and corn **(B)** microbiomes. Branches and labels of major lineages of bacteria from plant microbiomes are colored (pink = Bacteroidia, red = Bacilli, brown = Actinobacteria, green = alpha-proteobacteria, blue = beta-proteobacteria and purple = gamma-proteobacteria). The color strip around the outside of the tree represents ASV origin, with HTPC ASVs in orange, and microbiome ASVs in blue.

#### Recovered isolates from pea and corn roots

Based on the purity/number of reads we selected ASVs to attempt to culture over 100 unique ASVs from each collection. The criteria for selection of attempted recovery was ASVs with at least 20% purity and at least 60 reads (121/244 in pea; 200/250 in corn). Following streak-plate purification of the isolates, they were authenticated by full-length 16S rRNA gene sequencing. We successfully recovered ∼50% of the putatively culturable isolates (79/121 in pea; 136/200 in corn), with a slight bias toward success in the corn microbiome. We also noticed consistently that the number of colonies achieved during culture streaking approximately matched the relative number of reads for each isolate. The resulting isolates from pea and corn microbiomes showed similar phylogenetic distributions (**Figure 2**). Such a collection represents a powerful tool to contrast synthetic communities derived from different hosts^42^.

**Figure 2:**
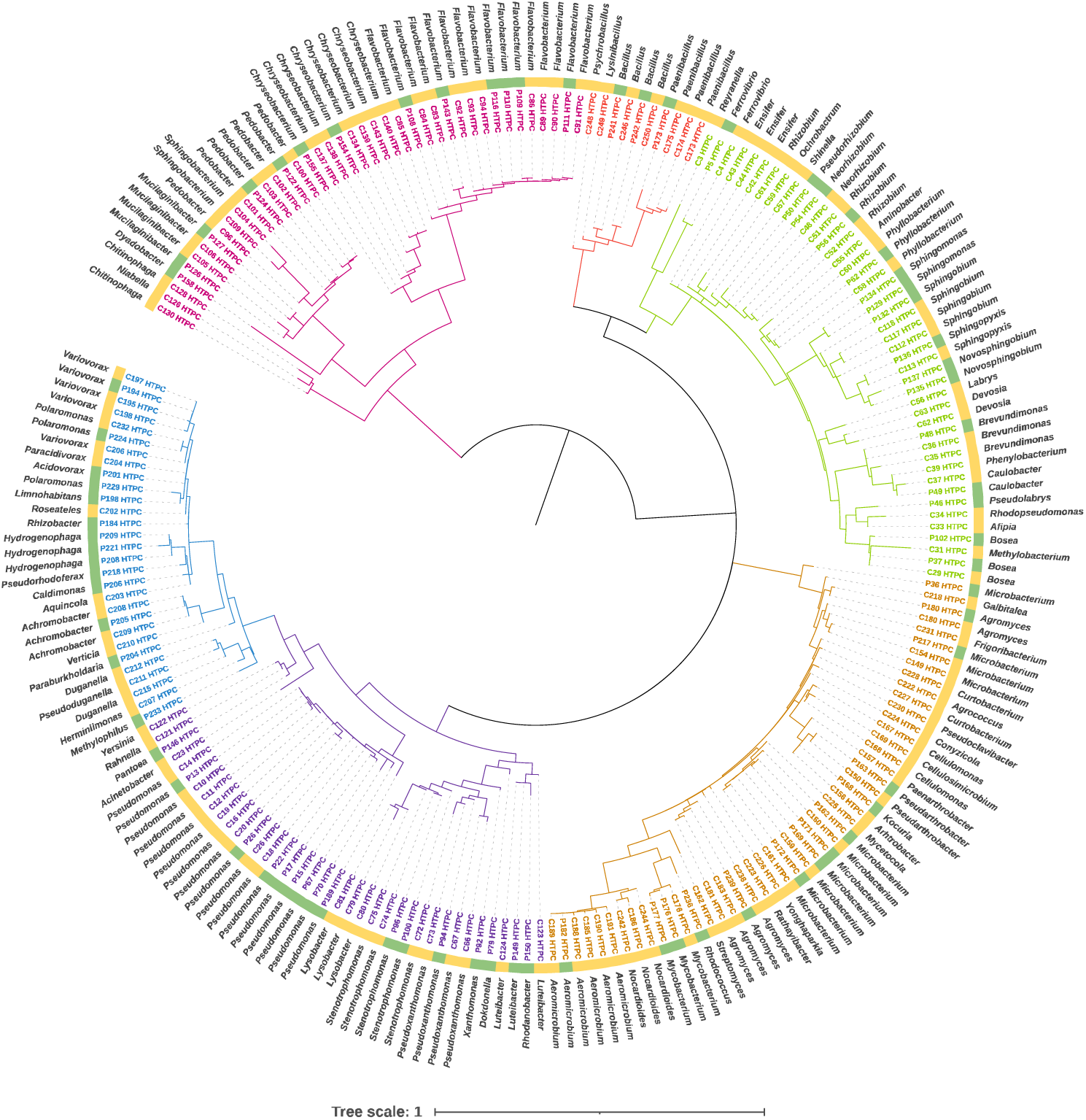
Maximum likelihood phylogenetic trees comparing the 215 cultured microbes from this work. Branches and labels of major lineages of bacteria from plant microbiomes are colored (pink = Bacteroidia, red = Bacilli, brown = Actinobacteria, green = alpha-proteobacteria, blue = beta-proteobacteria and purple = gamma-proteobacteria). The color strip around the outside of the tree represents ASV origin, with HTPC ASVs isolated from pea in green, and ASVs isolated from corn in yellow.

#### Comparison to other Methods

This method builds on advances in culturomics derived from the literature including colony picking, high-throughput cultivation and cell sorting^34,35,43,44^. In specific, we adapted and optimized a protocol for high-throughput culturomics from the plant root microbiome which we applied to field-grown crop plants from North Dakota, USA^34^. Based on our optimizations we suggest increasing the dilution series for field plant-root samples over those previously recommended^34^. To decrease the cost of identification primers and increase accuracy, we employed a method involving a two-step barcoding PCR. Compared to similar protocols, our protocol does not require gel cutting, which decreases the cost, time and labor. It can also be deployed using a high-throughput library preparation protocol utilizing an acoustic liquid handling robot we recently reported^38^. The described protocol employs the high throughput identification of bacteria by sequencing the V4 region of the 16s rRNA gene. The V4 region of the 16s rRNA gene was been pointed out as the most reliable^45^, reproducible^46^, informative^47^ and most similar to shotgun sequencing results^48^ compared to other regions. In terms of bioinformatics, this protocol uses a pipeline built around the DADA2 package^37^ in R^49^ to demultiplex the primers. DADA2 is one of the most used pipelines for processing amplicon sequencing data with great results and resolution ^50^. Moreover, due to the versatility and open-source characteristics of R, several packages to analyze microbiome data have been developed^51–55^. This protocol uses SILVA database^56^ for taxonomy assignment of the sequenced reads, the recommended database in the DADA2^37^ workflow. SILVA is the largest 16s rRNA gene dataset compared to RDP, Greengenes, NCBI and OTT ^57^.

### Conclusions

This study optimized and applied a high-throughput culturomics protocol to isolate and identify culturable bacterial taxa from the root microbiomes of field-grown pea and corn plants using NGS. The protocol combined dilution-to-extinction culturing with a two-step PCR and amplicon NGS strategy, successfully recovering around 50% of the unique putatively culturable amplicon ASVs identified through our unique bioinformatic pipeline. The resulting bacterial collection is taxonomically diverse, comprising plant-associated bacteria with potential applications in synthetic community design, evolutionary studies, comparative genomics, microbial ecology, and microbiome research. The optimized protocol demonstrated enhanced accuracy, cost-effectiveness, and high recovery rates, establishing a scalable and efficient method for large-scale culturing efforts in plant-microbiome research.

## 8. Author statements

### 8.1 Author contributions

Conceptualization, E.L.E., and B.A.G.; Methodology, E.L.E, N. D., K. M., and B.A.G.; Validation E. L. E., M. M., M. A. M., R. W.; Writing – original draft preparation, E. L. E., N. D., B.A.G.; Writing – review and editing, E. L. E., N. D.,M. M., M. A. M., R. W., B. A. G.; Supervision, B.A.G, E. L. E.

## 8.4 Acknowledgements

We thank Audrey Kalil for facilitating the sampling of diseased pea plants in Williston, ND. We also thank Nonoy Bandillo and Mike Ostlie for allowing us to sample pea salinity trials in Carrington, ND. We thank Kelsey Griesheim and Joel Bell for facilitating the sampling of a corn N-application trial in Gardner, ND. Scott Hoselton and Kaycie Schmidt in the Thomas Glass Biotech Innovation Core and Megan Ramsett in the Department of Microbiological Sciences, North Dakota State University contributed technical support for this work. This work used resources of the Center for Computationally Assisted Science and Technology (CCAST) at North Dakota State University, which were made possible in part by NSF MRI Award No. 2019077.

---

**Supplementary Table 1.**
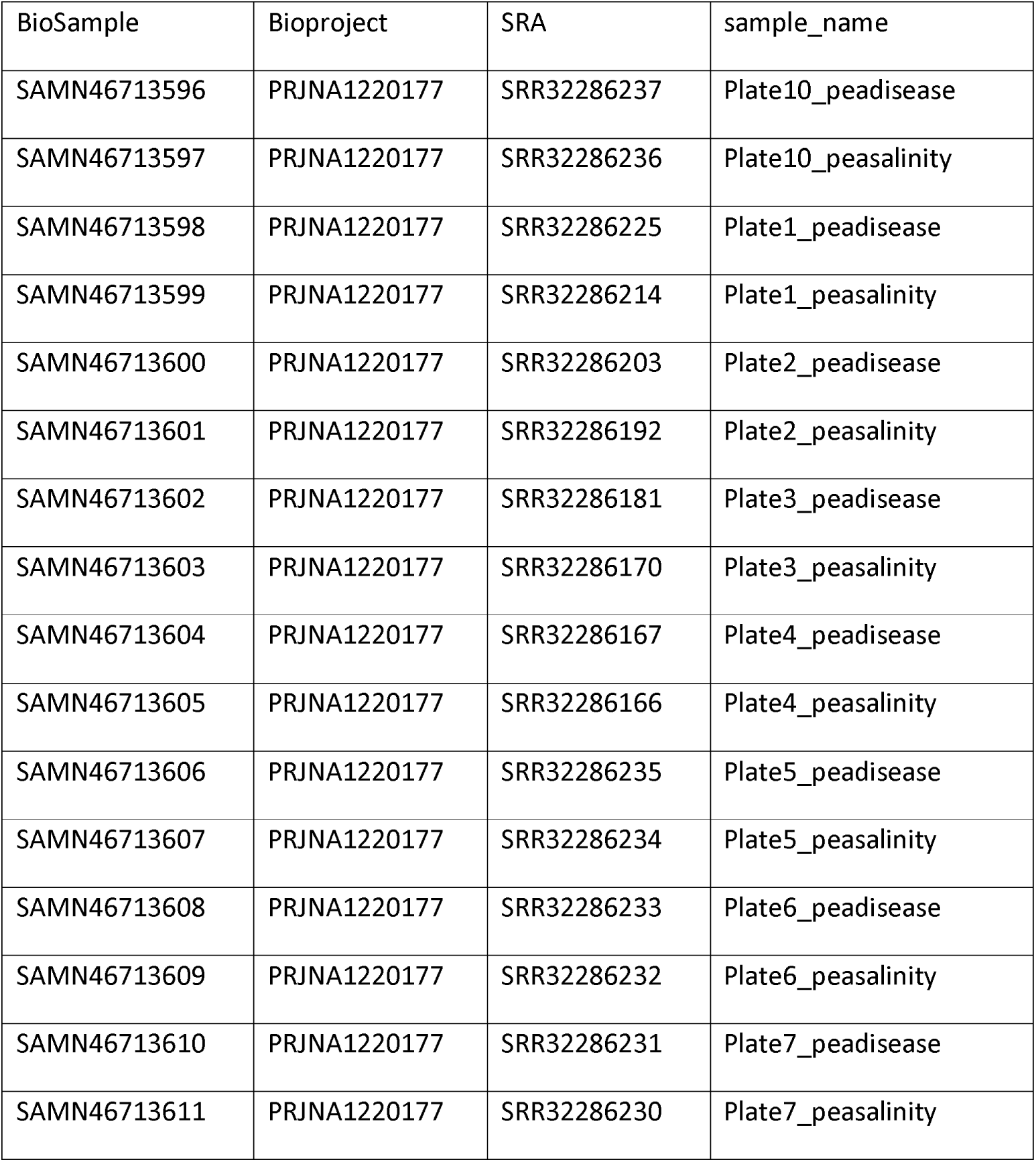
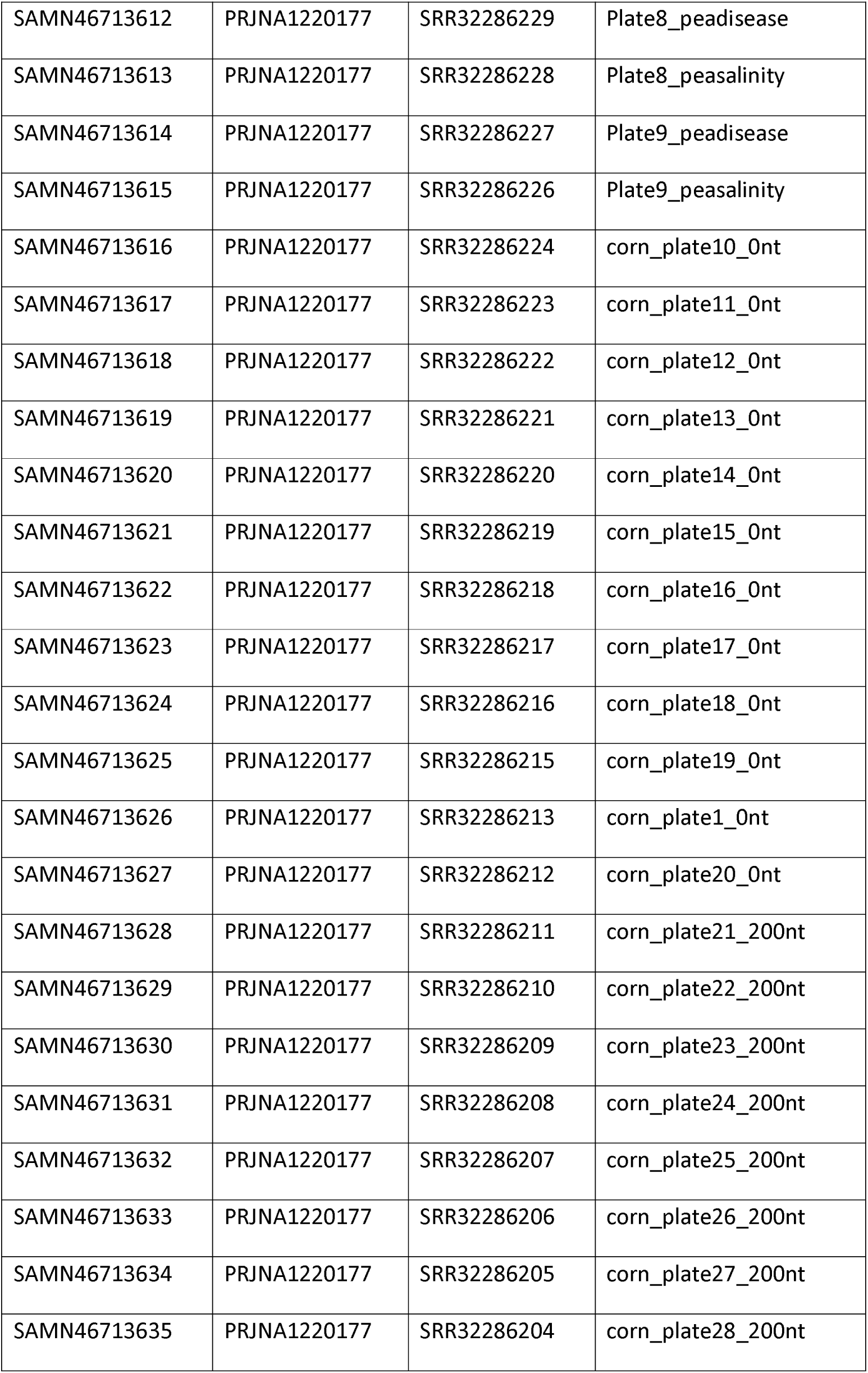
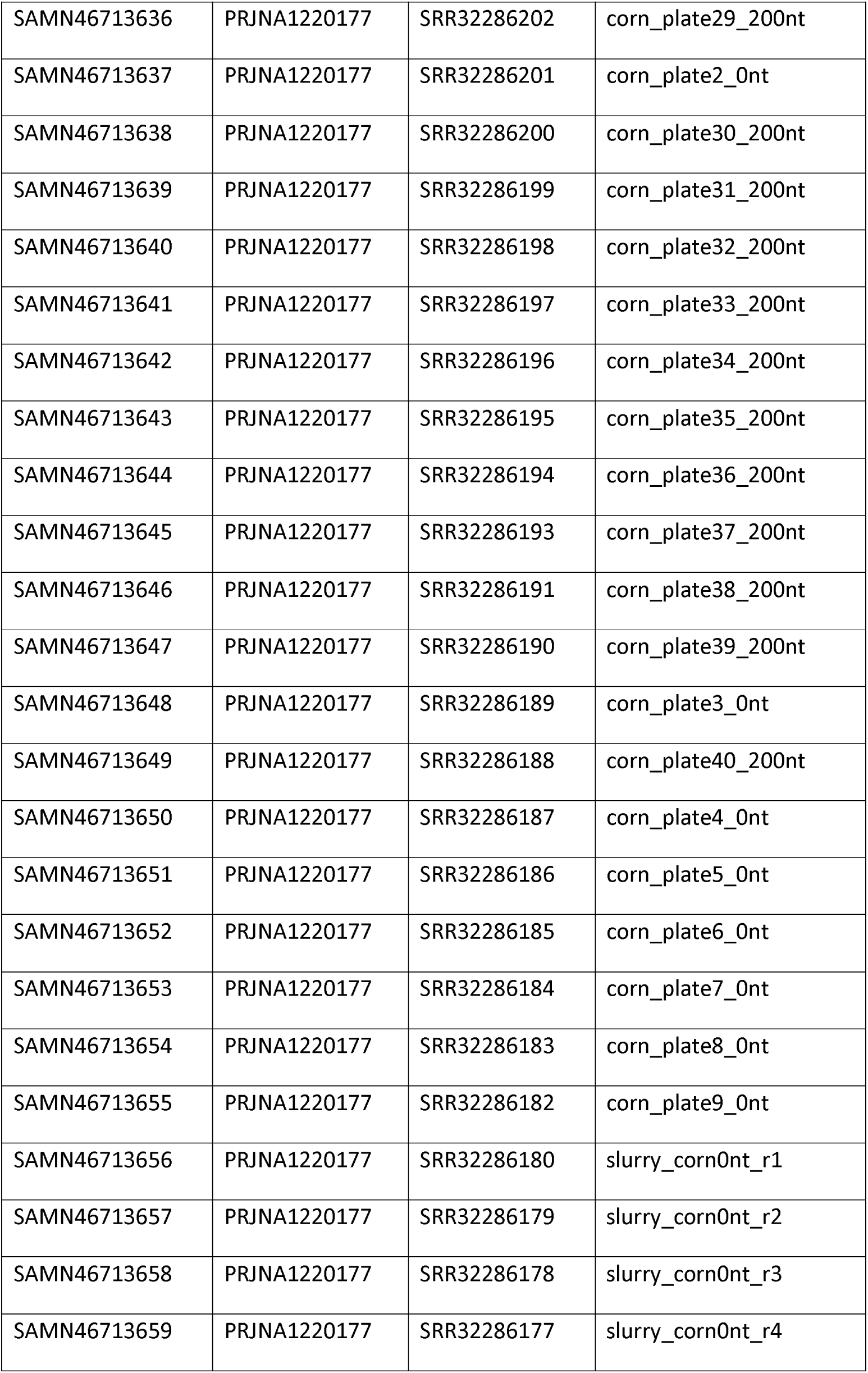
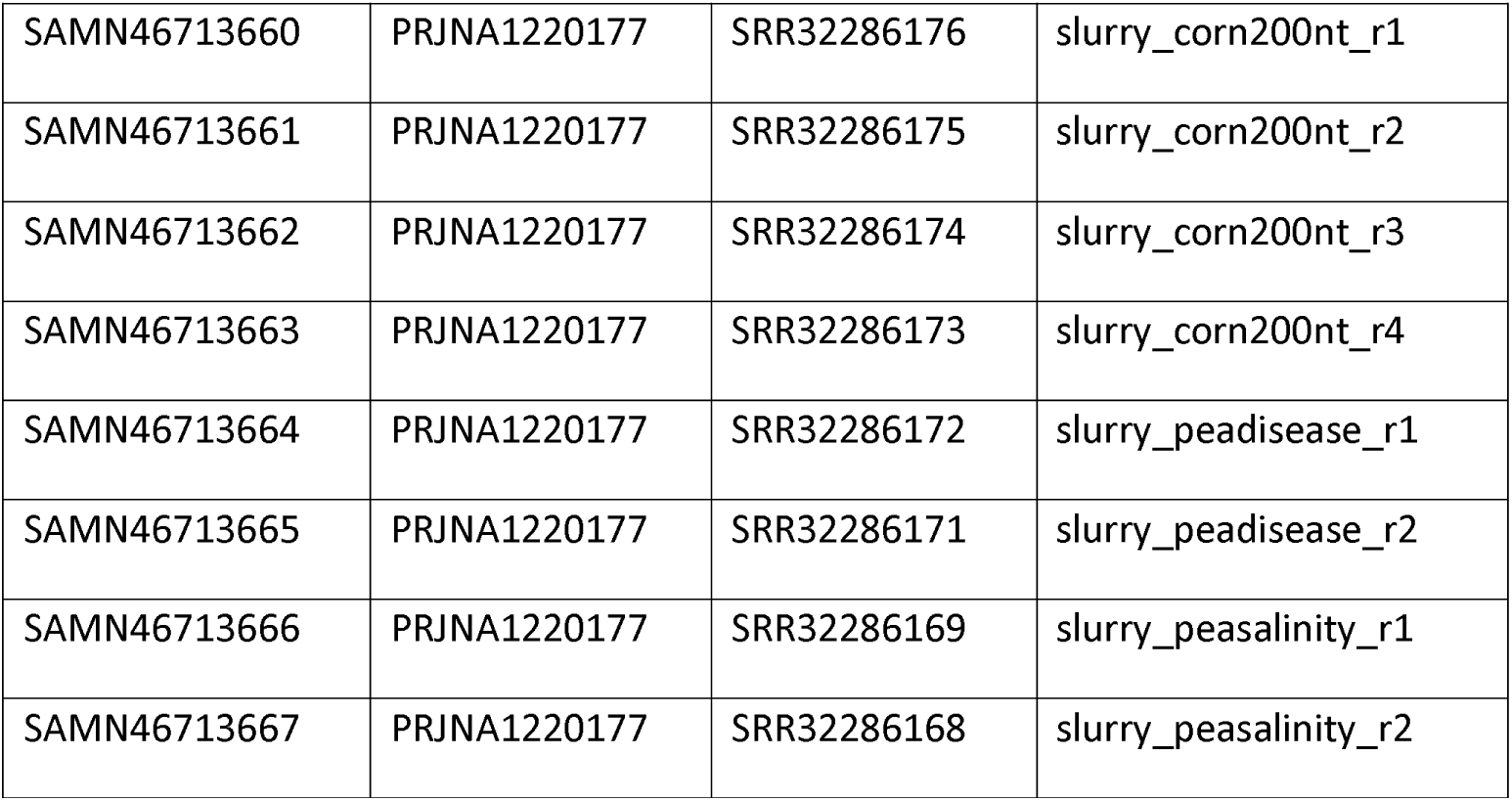
SRA accessions.

**Supplementary Table 2.**
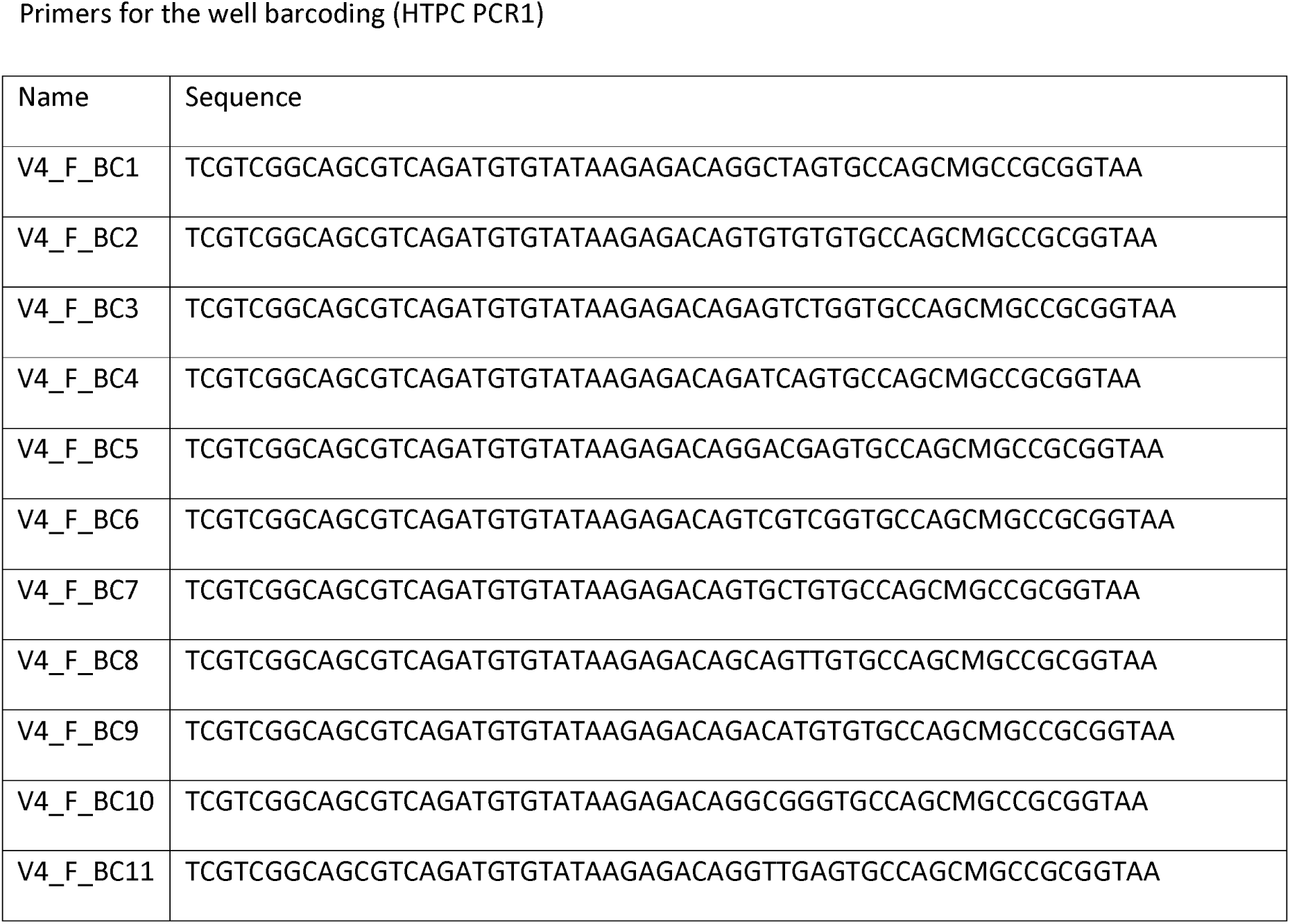
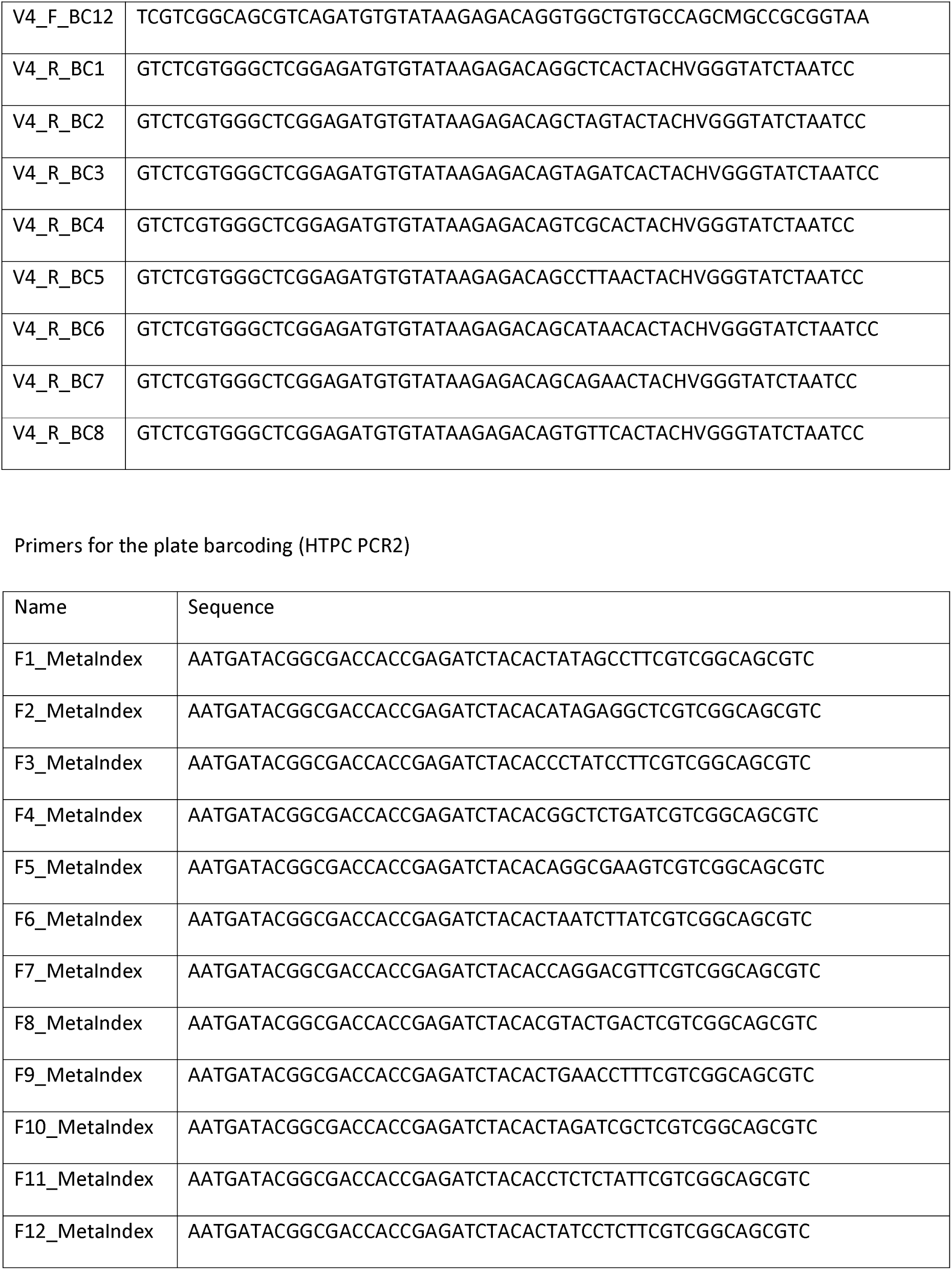
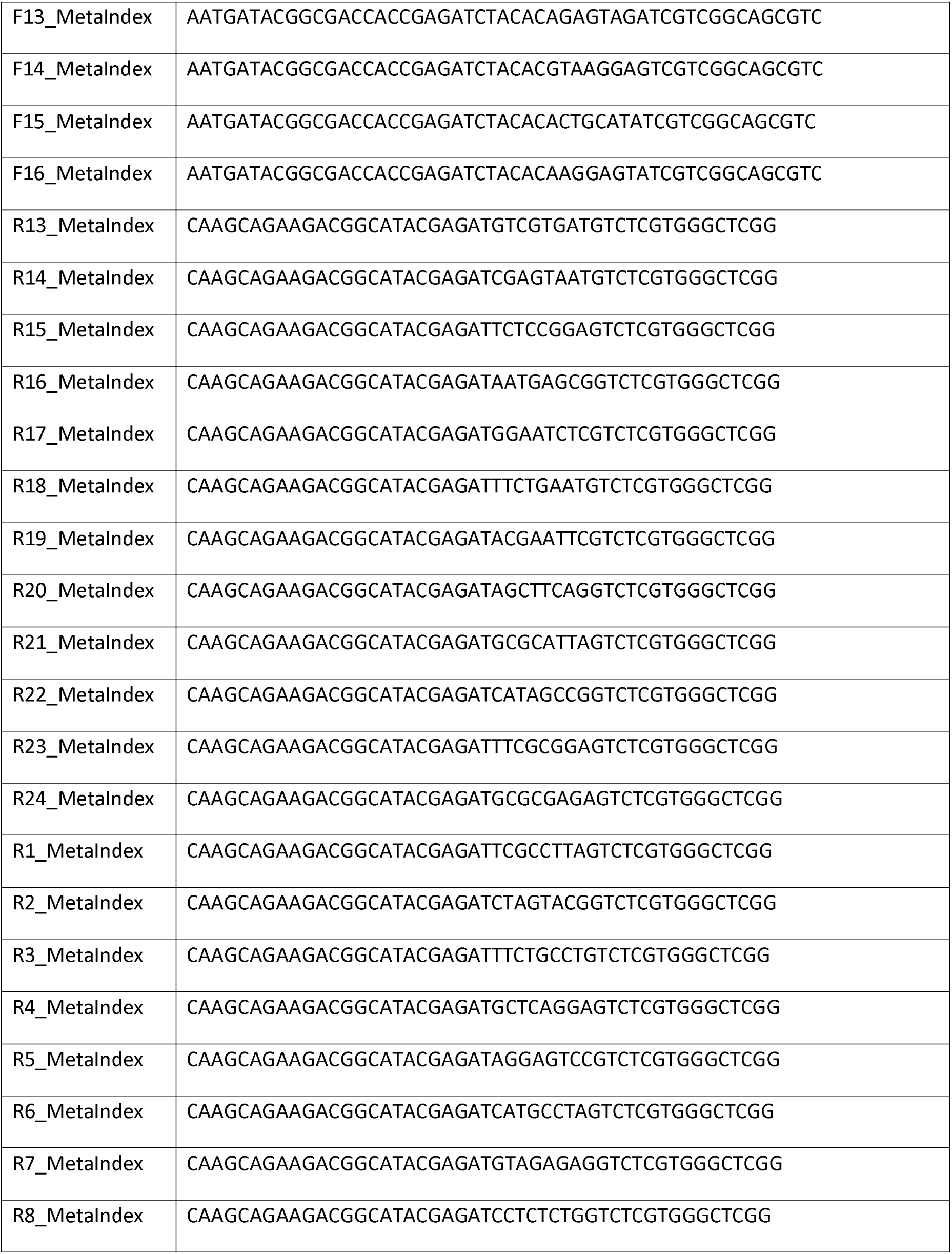
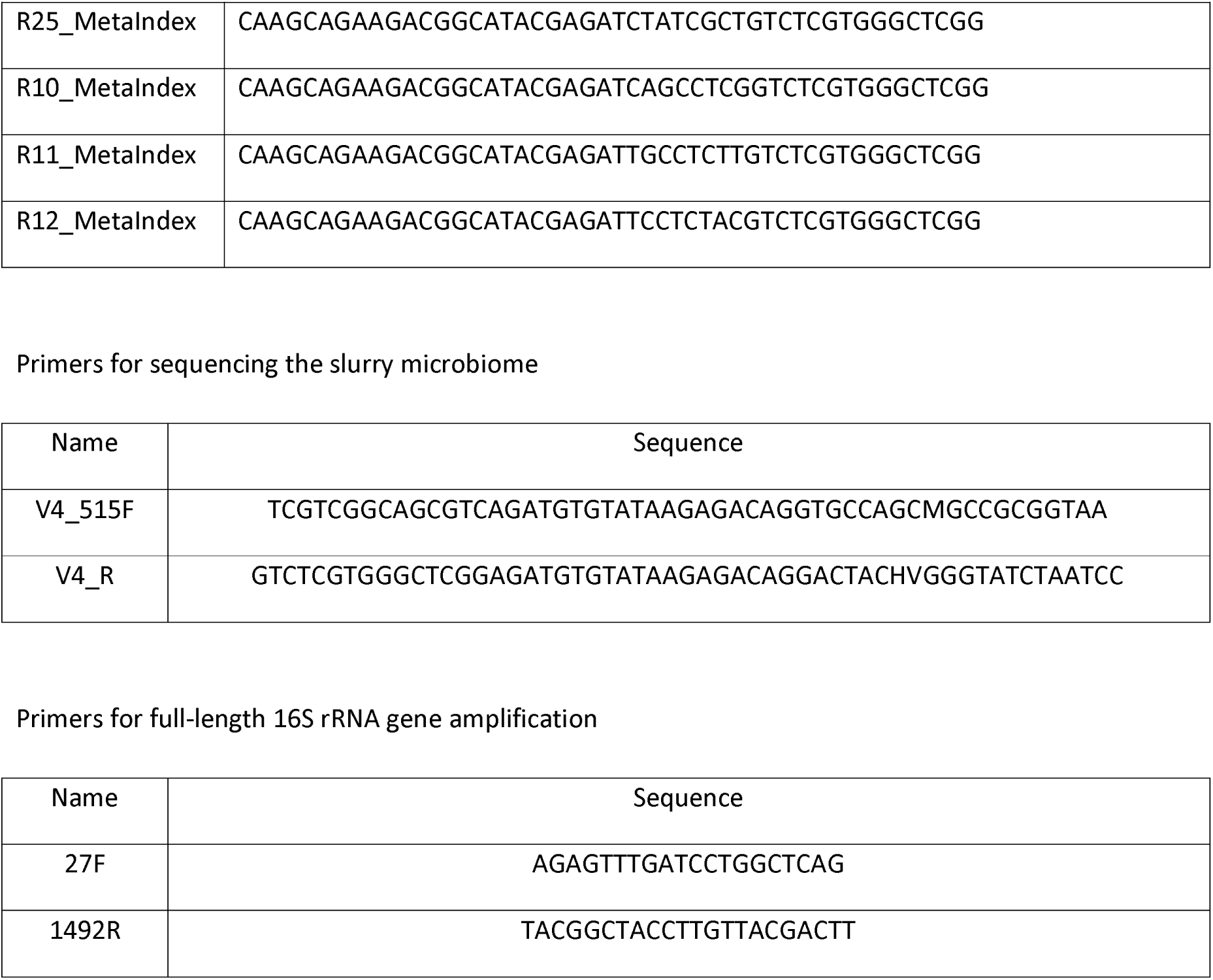
Primers used in this work.

